# Integration of graph neural networks and genome-scale metabolic models for predicting gene essentiality

**DOI:** 10.1101/2023.08.25.554757

**Authors:** Ramin Hasibi, Tom Michoel, Diego A. Oyarzún

**Affiliations:** Computational Biology Unit, Department of Informatics, University of Bergen, Norway; School of Biological Sciences, University of Edinburgh, UK; School of Informatics, University of Edinburgh, UK; The Alan Turing Institute, London, UK

## Abstract

Genome-scale metabolic models are powerful tools for understanding cellular physiology. Flux balance analysis (FBA), in particular, is a popular optimization approach for predicting metabolic phenotypes under genetic and environmental perturbations. In model microbes such as *Escherichia coli*, FBA has been successful at predicting essential genes, i.e. those genes that impair survival when deleted. A central assumption in this approach, however, is that both wild type and deletion strains optimize the same fitness objective. While the optimality assumption may hold for the wild type metabolic network, deletion strains are not subject to the same evolutionary pressures and knock-out mutants may steer their metabolism to meet other objectives for survival. Here, we present FlowGAT, a hybrid FBA-machine learning strategy for predicting essentiality directly from wild type metabolic phenotypes. The approach is based on graph-structured representation of metabolic fluxes predicted by FBA, where nodes correspond to enzymatic reactions and edges quantify the propagation of metabolite mass flow between a reaction and its neighbours. We integrate this information into a graph neural network that can be trained on knock-out fitness assay data. Comparisons across different model architectures reveal that FlowGAT predictions for *E. coli* are close to those of FBA for several growth conditions. This suggests that gene essentiality can be accurately predicted by exploiting the network structure of metabolism, without additional assumptions beyond optimality of the wild type. Our approach demonstrates the benefits of combining the mechanistic insights afforded by genome-scale models with the ability of deep learning models to extract patterns from complex data.

## I INTRODUCTION

The identification of essential genes is crucial for understanding the minimal functional modules required for survival of organisms [48]. Gene essentiality has key applications in biomedicine and biotechnology, for example to identify therapeutic targets in complex diseases [6], find strategies to combat pathogens [16], or optimize chemical production in genetically-engineered microbes [10]. Identification of essential genes requires screening assays where multiple knock-out mutants are phenotyped with a suitable fitness selection strategy. Such screens have been performed on many organisms, including model microbes such as *Escherichia coli* [2, 31, 37], *Saccharomyces cerevisiae* [45] and *Bacillus subtilis* [26], as well as pathogens such as *Candida albicans* [38] and *Aspergillus fumigatus* [23]. In human cells, recent work has produced high resolution deletion assays [6], leveraging progress in high-throughput technologies such as RNA interference and CRISPR-based screens [48] to produce detailed maps of gene essentiality in different conditions.

As a result of the cost and complexity of knock-out fitness assays, there is a growing interest in computational methods that can complement the experimental work with *in silico* prediction of fitness effects. These computational approaches often employ machine learning techniques combined with information from protein sequence, gene homologies, gene-function ontologies, and protein interaction networks [1, 8, 29, 30, 49]. In the case of metabolic genes, i.e. those that code for catalytic enzymes in metabolic pathways, Flux Balance Analysis (FBA) is a widely employed method for predicting essentiality [34]. There are numerous variants of FBA and its related algorithms [28], but at its core FBA computes genome-scale flux distributions that optimize a cellular fitness objective. Such objective is typically taken to be the cellular growth rate modelled as a linear combination of synthesis rates of amino acids, lipids and other biomass components. By imposing constraints on each metabolic flux, FBA problems can be solved with efficient linear programming algorithms, which allows to rapidly simulate the impact of gene deletions on the predicted growth rate and draw predictions on the essentiality of metabolic genes.

Flux Balance Analysis has shown good prediction accuracy for gene essentiality in the *E. coli* bacterium [31] and other model microbes, but predictions for eukaryotes and higher-order organisms have produced mixed results [18, 22]. The quality of FBA predictions have also been shown to vary strongly across published models as well as the performance metrics employed to quantify prediction accuracy [4]. An often overlooked limitation of the FBA approach is the tacit assumption that the metabolism of deletion strains optimizes the same objective as the wild type. In many cases, deletion strains display suboptimal growth phenotypes [44] and they are not subject to the same long-term evolutionary pressures as the wild type. It has also been postulated that deletions of metabolic genes can alter cell physiology to meet other objectives for survival; for example, an early work hypothesized that knockout strains may minimize their phenotypic deviation from the wild type [44], while various works have explored the impact of alternative objective functions [17, 42] and multiobjective optimization principles [43] in the classic FBA formulation.

Here, we sought to determine if gene essentiality can be predicted directly from wild type metabolic phenotypes. We developed a hybrid algorithm to predict gene essentiality using a combination of FBA and deep graph neural networks trained on knock-out fitness data. This approach does not require the assumption of optimality of deletion strains and takes maximal advantage of the inherent graph structure of cellular metabolism. Early attempts to augment the predictive power of FBA with machine learning explored the use of flux features for improved prediction of gene essentiality [33, 36], and other works have attempted to predict essentiality from the metabolic graph topology [15]. Most recently, several authors have developed integrated pipelines aimed at improving FBA predictions for biomedical [27, 35] and biotechnology tasks [13, 41].

In our approach, starting from wild type FBA solutions we first represent genome-scale flux distributions as a weighted digraph in a space of reaction nodes, and employ a flow-based representation for each node based on the redistribution of chemical mass flows between various paths in the graph. To integrate the graph structure and node features into a single predictive model, we employ a Graph Neural Network (GNN) with an attention mechanism [46] termed FlowGAT. We show that FlowGAT can be trained on a small amount of labelled data from knock-out screens. We demonstrate the effectiveness of this approach using the latest metabolic model of *E. coli*, and show that prediction accuracy near the FBA gold standard. Moreover, model predictions appear to generalize well across various growth conditions without the need for further training data. The results highlight the advantages of integrating FBA pipelines with state-of-the-art machine learning algorithms for improved phenotypic predictions.

## II RESULTS

### A Model architecture and training

In this paper, we propose FlowGAT, a GNN based model to predict gene essentiality from graphs generated from FBA solutions. As shown in Figure 1A, each node in the graph corresponds to a metabolic reaction, and we pair each node with a set of flow-based features and binary essentiality labels obtained from knock-out fitness assays. The graph structure and node features are integrated into a GNN for binary classification, so as to use a message passing scheme to propagate node features through the structure of the graph; this allows learning a rich embedding of the input that contains information from the *k*-hop neighbourhood of each node [19]. We next detail the different components of the model and our training strategy.

**FIG. 1.**
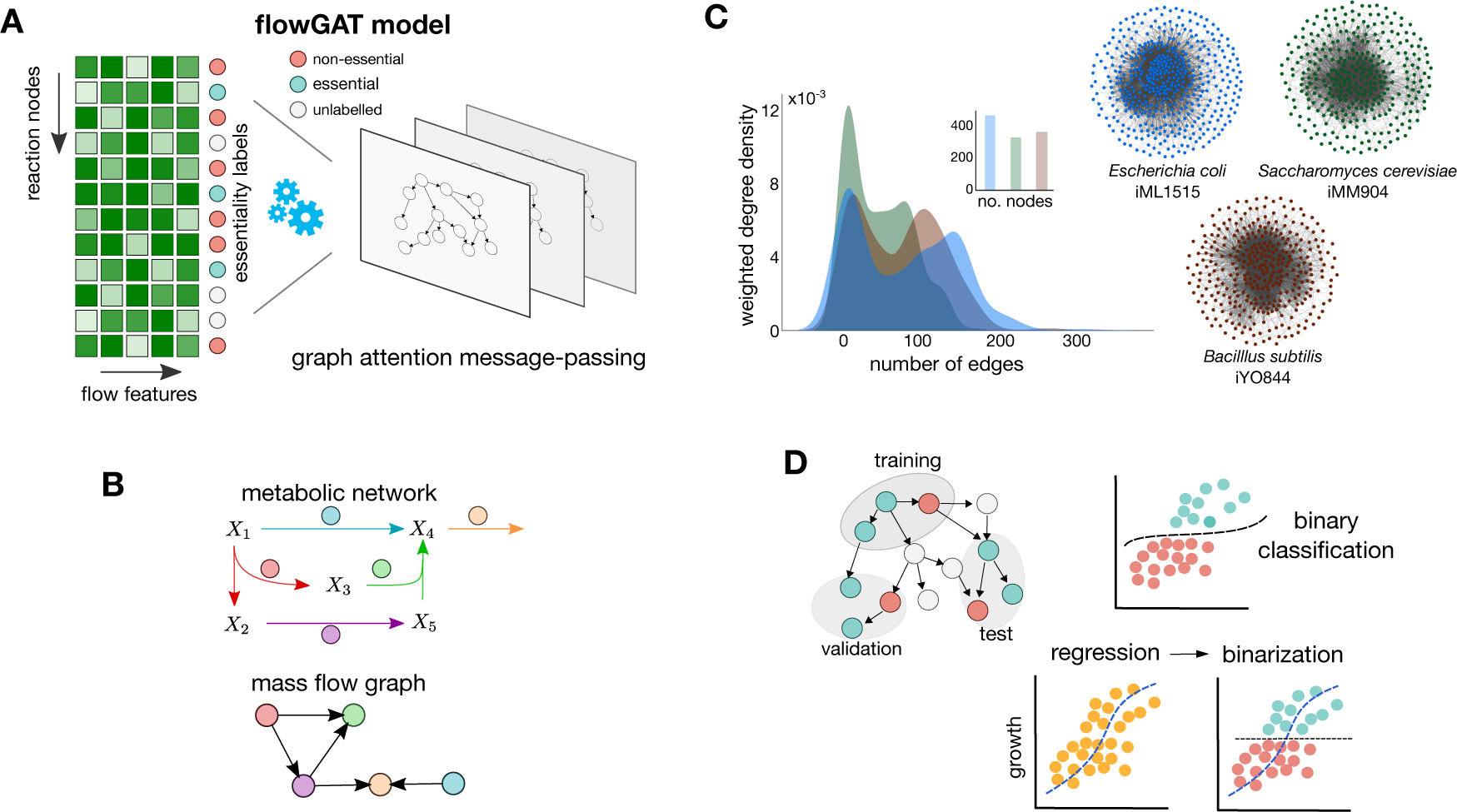
Elements of the FlowGAT model for gene essentiality prediction. (**A**) Schematic of the FlowGAT architecture proposed in this paper. The model integrates a digraph representation of FBA solutions (Mass Flow Graphs, MFG), where nodes are reactions and edges encode the re-distribution of metabolite mass flows between reactions. We featurize each node with flow-based scores and label them as essential or non-essential using data from gene knock-out assays. Using a graph neural network with an attention layer, FlowGAT predicts essentiality for unlabelled reactions. (**B**) Construction of mass flow graphs from FBA solutions. The top network is an exemplar metabolic network, and the bottom digraph is the corresponding MFG constructed; nodes are reactions and two nodes are connected if they share metabolites as reactants or products. The edge weights are computed from the metabolite mass flows as described in (3); more details on the MFG construction can be found in [3]. (**C**) Exemplar MFGs for several microbes computed from their genome-scale models using standard FBA [12] with the default growth condition in each case; density plots show the distribution of edge weights in each case. (**D**) For model training and validation, labeled nodes in the MFG are separated into training, validation and test sets. The validation set is used for early stopping and performance metrics are computed on the test set. We explored two training frameworks for FlowGAT; as a binary classifier and as a regressor of growth rate that can be binarized to produce essentiality predictions.

#### a. Graph construction

We consider metabolic networks with *m* metabolites and *n* enzymatic reactions described by the following differential equation model

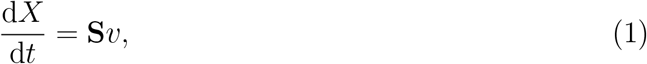

where *X* is an *m*-dimensional vector of metabolite concentrations, *v* is a *n*-dimensional vector of reaction fluxes, and **S** is a *n×m* stoichiometric matrix. In steady state, the relation **S***v* = 0 describes all flux vectors that can sustain a specific metabolic state. A common strategy to estimate *v* at the genome-scale is to employ FBA to compute a flux vector *v*^⋆^ that optimizes a meaningful biological objective; details on FBA can be found in the Methods section. To convert such FBA solution vectors *v*^⋆^ into a graph, we used the Mass Flow Graph (MFG) construction proposed by Beguerisse-Diaz *et al* [3] and illustrated in Figure 1B. Starting from the stoichometric matrix **S**, we first build a directed graph with reactions as nodes, where two nodes are connected if and only if the source reaction produces a metabolite that is consumed by the target reaction. Each edge in the graph has a weight *w*_*i,j*_ that represents the normalized mass flow from node *i* to node *j*. We first compute the flow of metabolite *X*_*k*_ from reaction *i* to *j* according to:

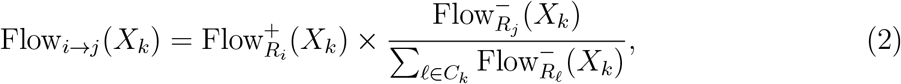

where 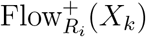 and 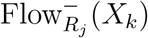 are the production and consumption flows of metabolite *X*_*k*_ by reaction *R*_*i*_, respectively. The set *C*_*k*_ contains the indices of all reactions that consume metabolite *X*_*k*_. The edge weight *w*_*i,j*_ is thus defined as the total mass flow between two nodes, aggregated over all metabolites *X*_*k*_ that are produced by node *i* and consumed by node *j*:

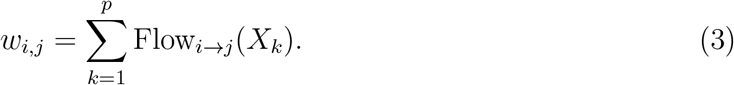

Mass flow graphs allow converting FBA solutions into a directed graph, and thus can be used to represent the network structure of metabolism in different growth conditions or genetic perturbations. In Figure 1C, we show MFGs built from genome-scale metabolic models for three model microbes available in the BiGG model database [24] (*Escherichia coli, Saccharomyces cerevisiae* and *Bacillus subtilis*). Further details on the construction of the mass flow graphs can be found in the Methods section.

#### b. Design of node features

Besides the graph topology, we ascribe a feature vector to each reaction node that can be exploited for improved performance by the representation learning approach. This approach is analogous to the structural and positional encoding schemes employed in graph Transformer architectures to feed models with extra information about the local connectivity of nodes [32]. Since the edge weights in (3) relate to the mass flow between reactions, we opted to employ flow-based features that can aggregate information on incoming and outgoing mass flows from each node. To this end, we employ the Flow Profile Encoding (FPE) first defined by Cooper and Barahona for general directed graphs [9]. Given a directed MFG with weighted adjacency matrix **A**, for each node *i* we define the inflow profile of length *k* as

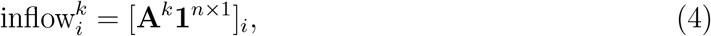

where **A**^*k*^ is the matrix *k*-th power, **1**^*n×*1^ is an *m*-dimensional vector of ones, and [*·*]_*i*_ is the *i*-th element of a vector. The inflow of node *i* is thus defined as the weight sum across all incoming paths of length *k*. We similarly define the outflow of node *i* as:

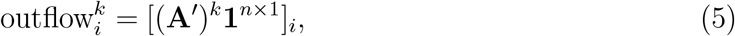

where **A**^*′*^ is the matrix transpose. We concatenate inflows and outflows up to maximal length *k*_m_ for each node:

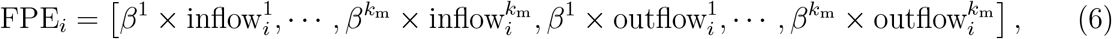

where *k*_m_ is a hyperparameter that defines the maximum path length, *β* = *α/λ*_1_ is a scaling factor, *λ*_1_ is the largest eigenvalue of the adjacency matrix **A**, and *α* is a hyperparameter that controls for the variable weights of the short and long paths; normalization by the largest eigenvalue *λ*_1_ ensures convergence for large *k*, in the sense that lim_*k*→∞_ ||**A**^*k*+1^||*/*||**A**^*k*^|| = *λ*_1_. The definition in (6) allows computing a feature vector of length 2*k*_m_ for each node in the graph.

#### c. Representation learning

Graph representation learning is concerned with mapping the nodes into a low dimensional vector which is optimized for downstream tasks such as classification or regression [21]. GNNs are a family of deep learning methods on graphs which obtain the embedding vector by incorporating the features of the node and its local neighbourhood according to a customized message passing scheme called Message Passing Neural Network (MPNN) [19]. Doing so helps the model capture local and global structural information about the graph and results in a more expressive embedding space. For more information about MPNNs refer to the Methods section. In this study, we employ a MPNN architecture named Graph Attention (GAT) to compute the neighbourhood information importance in finding the representation of the node [46]. Each layer *l* of GAT updates the representation of node *i* according to:

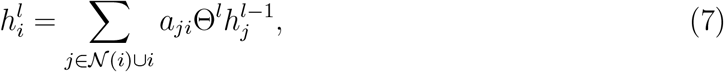

where 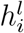 is the representation vector, 𝒩 (*i*) is the set of neighbouring nodes for node *i*, Θ is a set of differentiable weights, and *a*_*ij*_ is an attention coefficient that is dynamically calculated for each node *j* ∈ 𝒩 (*i*) ∪ *i* as

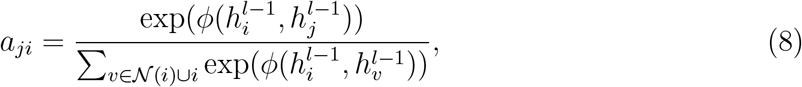

where *ϕ* is a differentiable function optimized through gradient descent optimization algorithms [39]. Details on the message passing and attention schemes can be found in the Methods section.

#### d. Data pre-processing

FlowGAT can be trained on knock-out growth assay data, where each gene is labelled as non-essential (0) or essential (1) depending on whether a fitness score is above or below prescribed threshold. For model training, the binary gene labels must be converted into their corresponding reaction node labels in the MFG. To this end, we use the Gene-Protein-Reaction (GPR) map included in genome-scale metabolic models. The GPR is a Boolean function that specifies which gene codes for which proteins, and conversely how each protein affects a metabolic reaction. The GPR can account for reactions that are catalyzed by multiple enzymes or by enzymatic complexes encoded in multiple genes. For those genes that map one-to-one into a single reaction, we transferred the gene label directly into a reaction label. For those genes that map into multiple reactions (many-to-one), we transferred the gene label to all reactions deactivated by the gene deletion. When multiple genes map to multiple reactions (many-to-many), the structure of the GPR does not allow to infer reaction labels from gene labels, and therefore we considered such reactions as unlabelled. Note that the data also contains nodes that lack essentiality labels because their corresponding genes have not been measured in the growth assay. The unlabeled nodes are made available for model training to make sure that the graph representation learning can take advantage of the full graph structure without limiting the representation power of the GAT; the classification loss for training and evaluation of the model is only calculated on the labeled nodes. We also note that the reaction labels are typically imbalanced because the MFG is enriched for essential reaction nodes. By definition in (3), those reactions with zero flux in the wild type FBA solution will have nil edge weights and thus are disconnected from the graph. During training, the model has access to the features of all nodes (labeled and unlabeled) through the message passing, but the training loss is calculated on the labels of the training nodes in semi-supervised fashion (Figure 1D). Details on model training can be found in the Methods section.

### B Performance evaluation of FlowGAT

To evaluate the performance of FlowGAT, we employed the growth knock-out data for the *Escherichia coli* bacterium reported by Monk and colleagues [31], and the iML1515 genome-scale model reported in the same work. We chose the *E. coli* model because it is the most complete and best curated metabolic reconstruction in the literature, and thus allows us to mitigate the impact of misclassification errors caused by poor model quality and focus on the predictive power of FlowGAT itself. The dataset contains growth rate data for 3,892 *E. coli* genes grown in various carbon sources.

We built the MFG for *E. coli* using the wild type FBA solution using glucose as the sole carbon source and the default objective function included in the iML1515 model (growth rate). The resulting MFG has 444 nodes and after converting the gene labels to reaction labels with the GPR map we obtained 255 labeled nodes (191 essential, 64 nonessential). We first compared FlowGAT trained on binary cross-entropy loss with classical binary classifiers including Support Vector Classifier (SVC), Multi Layer Preceptron (MLP) classifier, and random forests (RF) classifier using the flow profile embeddings in (6) as feature vectors; details on model training and hyperparameter selection can be found in Methods. The results in Figure 2A show precision-recall curves, averaged across *N* = 50 rounds of training and testing (5 test folds with 20% of nodes resampled 10 times for model retraining); details on our strategy for model evaluation can be found in the Methods. Among the considered classifiers, FlowGAT achieves the best Area Under the Precision-Recall Curve (PRAUC) across all test folds and performs above the no-skill classifier while the classic models significantly underperform; we note that due to the class imbalance the baseline precision of the no-skill classifier is 74.9%.

**FIG. 2.**
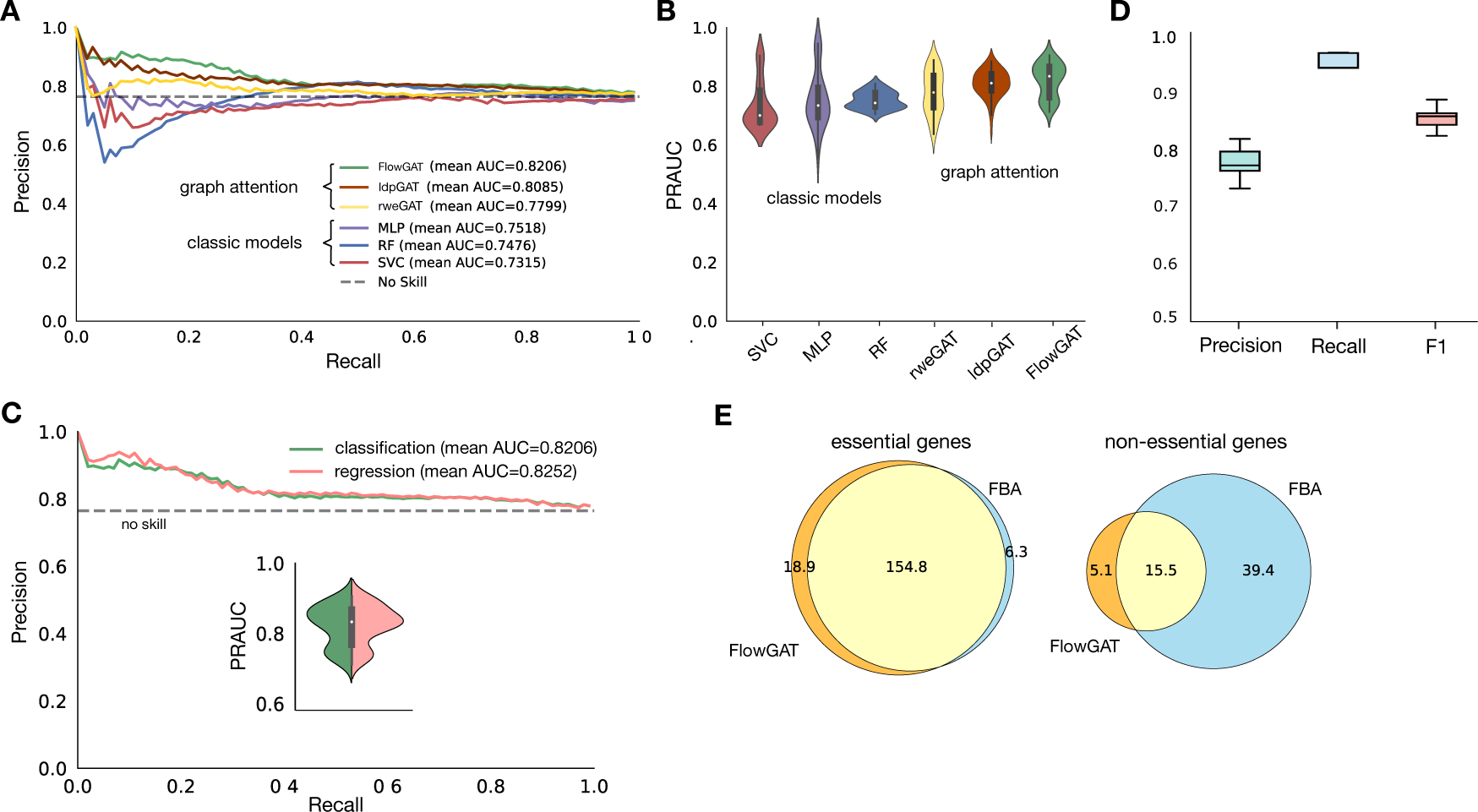
Performance of FlowGAT as predictor of metabolic gene essentiality in *Escherichia coli*. (**A**) Precision-Recall (PR) curves for classic binary classifiers and graph neural networks trained on essentiality measurements for *E. coli* growing in aerobic conditions with glucose as the sole carbon course [31]. Classic models were trained using the Flow Profile Embeddings (FPE) defined in Eq. (6) computed from wild type FBA solutions. The graph neural networks were trained using the mass flow graph and the FPE as node feature vectors, as well as two other popular node embedding techniques (Local Degree Profile, LDP [7] and Random Walk Embedding, RWE [11]). Results show PR curves averaged across 50 model evaluations consisting of 5 rounds of testing on 20% test folds and and 10 rounds of model re-training for different random seeds; the dashed line represented the precision (74.9%) of the no-skill classifier given the class imbalance of the data. (**B**) Distribution of PRAUC scores across the 50 evaluations. The graph attention models outperform classic binary classifier; among the three considered node embeddings, FPE provides the best performance. (**C**) Retraining FlowGAT as a regressor provides slight gains in performance; inset shows the distribution of PRAUC scores across 50 evaluations; the regressor was trained on growth rate data [31] and predictions are subsequently binarized to produce essentiality labels. (**D**) Exemplar classification results by the best performing model (FlowGAT trained as regressor, as in panel C); results show precision, recall and F1-score for the 50 evaluations and fixed classification threshold. (**E**) Comparison between FlowGAT and FBA predictions over the entire gene set in glucose MFG; Venn diagrams show the number of genes called correctly for each model, averaged across 10 rounds of re-training for different random seeds.

We also compared FlowGAT trained on two other popular node embedding techniques (Local Degree Profile (LDP) [7] and Random Walk Embedding (RWE) [11]) that have shown good performance in a number of tasks on molecular graphs, as well as two other message passing schemes (Graph Convolution Network [25] and Graph SAGE, see Supplementary Figure S1). Details on these additional node embeddings and message passing strategies can be found in the Methods section. The results (Figure 2A) show that graph attention delivers the best performance, and models trained on LDP and RWE node features are outperformed by the flow profile encodings, possibly because the former do not take account for directionality and weight of the edges of the MFG.

We further investigated the sensitivity of FlowGAT to the random seed employed for weight initialization; the distributions in Figure 2B show the PRAUC scores for all models across the 50 runs. The results suggest that FlowGAT performance is relatively robust; only the RF classifier delivers tighter predictions, but at the cost of an average performance below the no-skill baseline.

We finally sought to explore an alternative training scheme using a regression approach. Since the gene essentiality labels are based on a binarization of continuous measurements of growth rate, we reasoned that recasting the prediction problem as a regression task could improve performance. To this end, we employed the non-binarized fitness measurements of growth rate in Monk et al [31] and re-trained FlowGAT as a regressor using Mean Squared Error (MSE) loss on predicting the non-binary growth rate values; all model hyperparameters were left unchanged. Following the same evaluation scheme as the above, we used the FlowGAT regressor to predict growth rates for the reaction nodes in each test fold. We then used the predicted growth rate as classification scores and computed the precision-recall curve on the test fold. Upon comparison with the classification approach in Figure 2A–B, the regression results in Figure 2C led to a performance increase in terms of average PRAUC, as well as tighter predictions that are less sensitive to weight initialization.

After finding the best setting for FlowGAT in terms of architectural design choices, we fixed the cut-off threshold for the output prediction of FlowGAT trained as a regressor in Figure 2C to produce binary essentiality predictions for all 50 evaluations by classifying the nodes that score above the threshold as essential and others as non-essential. We measure the performance of FlowGAT in terms of three metrics of Precision, Recall, and F1 for the binary predictions (Figure 2D). The predictions of the FlowGAT regressor manage to keep both precision and recall above 75% and 90%, respectively.

To better understand the performance of our model, we compared the output of FlowGAT trained as a regressor (Figure 2C) with those from FBA applied to the genes that appear in the MFG. First, we mapped each reaction node back to its corresponding into gene labels using the GPR map. In total, 240 labeled genes appear in the constructed MFG (180 essentials and 60 non-essential. This number is lower than the number of reaction nodes (255) because some reactions correspond to the same gene based on many-to-one mapping that was used to assign labels to nodes earlier; for such genes, we aggregated reactions by maximum prediction value of corresponding reactions. We collected predictions across all genes and compared these results with the essentiality prediction of FBA for each gene in Figure 2E. The results suggest that both FlowGAT and FBA find most of the essential genes, but FlowGAT finds on average 19 essential genes that are misclassified by FBA. In the case of non-essential genes, however, we found that FlowGAT underperforms and misses more genes than FBA, likely as a result of non-essential genes being the minority class.

### C Essentiality prediction in different growth conditions

The essentiality of metabolic genes can be highly dependent on environmental conditions. Difference carbon sources can produce important differences on the metabolic phenotype and, as a result, some genes that are essential in one condition source may be non-essential in another one.

To test the predictive power of FlowGAT in other growth conditions beyond glucose, we trained the model using *E. coli* knock-out fitness data in ten other carbon sources that cover different entry points into central carbon metabolism [31]. We built the corresponding mass flow graphs from wild type FBA solutions of the iML1515 model instanced to each carbon source. To build condition-dependent graphs, we constrained the flux of each nutrient exchange reaction to a fixed value (Supplementary Table S2). This resulted in 10 different MFGs that differ on their nodes and their edge weights. Inspection of the reaction nodes per graph (Figure 3A) reveals differences across graphs for reactions that become active for specific carbon sources, as well as a large number of reactions that are shared across conditions. We then evaluated the performance of FlowGAT trained in each of the ten mass flow graphs, using the growth knock-out fitness data and the regression strategy of Figure 2C; model hyperparameters were left unchanged. In each graph, the number of essential and non-essential nodes varies and thus, the no-skill baseline varies depending on the class imbalance in that graph. As seen in Figure 3B, we found that while the PRAUC scores vary across growth conditions, in all cases FlowGAT outperformed the no-skill classifier by at least 6%. These encouraging results can likely still be improved by introducing condition-specific hyperparameters for the FlowGAT architecture.

**FIG. 3.**
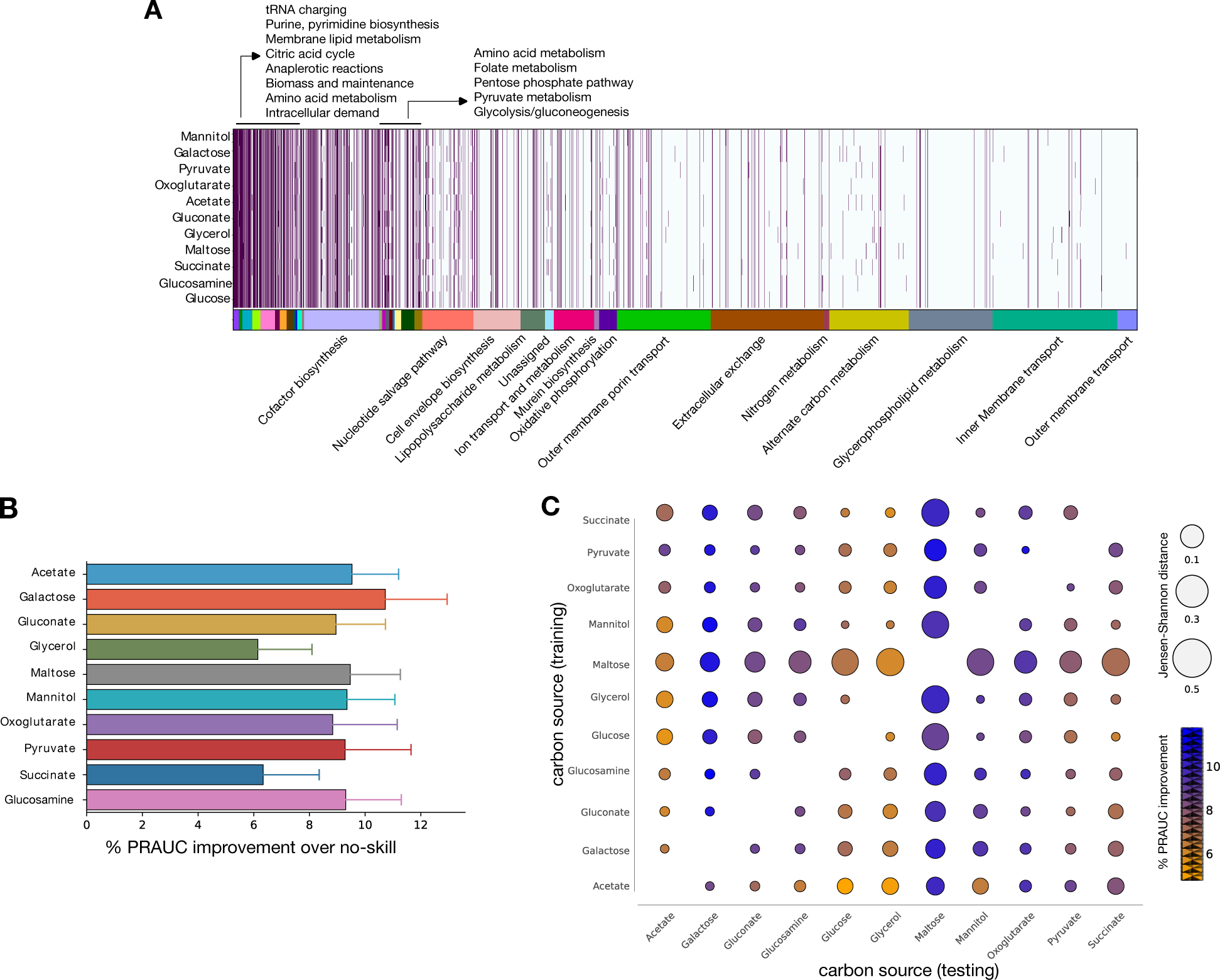
Essentiality predictions of FlowGAT for *Escherichia coli* growing in different carbon sources. **(A)** Heatmap of reactions present in mass flow graphs (MFG) computed from wild type FBA solutions computed for ten carbon sources. Each MFG is obtained through changing the carbon source; the color bar denotes the different metabolic subsystems as annotated in the latest genome-scale metabolic model iML1515 [31]. **(B)** Prediction performance of FlowGAT trained and tested on the different condition-dependent MFGs. Bars show the average improvement of PRAUC scores across the 50 evaluations with respect to the no-skill classifier; error bars denote one standard deviation of the PRAUC. Full precision-recall curves for each case can be found in Supplementary Figure S2. **(C)** Performance of FlowGAT in cross-testing across different carbon sources; in each case, the model was trained on one graph and tested on the nodes of all other graphs, totalling 90 cross-test evaluations. The color bar indicates the improvement in PRAUC over the no-skill classifier; the bubble radius denotes the graph-to-graph distance computed as the Jansen-Shannon divergence between the distribution of node features.

We finally aimed to determine the ability of FlowGAT to generalize predictions across growth conditions. We conducted a cross-training evaluation, where the model was trained on a mass flow graph and fitness data from a single carbon source, and tested on the reaction nodes in a different growth condition. To this end, we also included the FlowGAT model for glucose discussed in the previous section. As shown in Figure 3C, all 90 cross-tests show an improvement in PRAUC with respect to each MFG no-skill classifier. Although each FlowGAT model was trained on a different graph and fitness data, these results suggest that the model captures a well-performing representation of the data. To test if this is a result of the similarity between the nodes present in each graph (Figure 3A), we quantified the graph-to-graph similarity using the distance between the distribution of node features. We estimated the probability density function of the flow profile encodings for each graph using kernel density estimation and computed the Jensen Shannon divergence between all pairs of distributions. The results (Figure 3C) do not show a correlation between graph similarity and the PRAUC scores. For example, the maltose graph embedding is nearly equidistant from both the acetate and galactose graphs, but FlowGAT trained on maltose has a performance of approximately 5% better when tested on galactose than in acetate. Likewise, FlowGAT trained on mannitol performs better when tested in galactose than glycerol, despite the galactose graph being more dissimilar to the mannitol graph. These observations suggest that the generalization performance of FlowGAT results from its representation power rather than the similarity between the input graphs.

## III DISCUSSION

Gene essentiality refers to the concept that some genes are indispensable for the survival of an organism. These genes often encode proteins that play critical roles in fundamental cellular processes needed for growth. Since the quantification of gene essentiality requires knock-out fitness assays across a large number of genes and growth conditions, there is substantial interest in computational methods that can aid the identification of genes from a reduced number of measurements. In this paper, we presented FlowGAT, a graph neural network that can be trained on knock-out fitness data to predict the essentiality of metabolic genes. The architecture exploits the inherent graph structure of metabolic fluxes predicted by Flux Balance Analysis through a combination of mass flow graphs and node features that describe local connectivity.

Using data from *E. coli* and its latest genome-scale metabolic model, we show that FlowGAT can identify most of the genes that are correctly called as essential by Flux Balance Analysis, and even correct some of its misclassified essential genes. Our approach is based solely on the wild type phenotype predicted by FBA; since it does not require the assumption of optimality in the deletion strains, FlowGAT may provide benefits when applied to organisms where the growth optimality assumption is not warranted. Additionally, FlowGAT displays encouraging generalization power across growth conditions, even in cases where the underlying graphs and node features differ substantially. This observation suggests that the proposed architecture and feature extraction method can learn internal representations that are useful predictors of gene essentiality. We also found, however, that FlowGAT struggled to predict non-essential genes and can be substantially outperformed by traditional FBA. This phenomenon could arise from various sources, such as the data imbalance that in our case favors essential labels, or because predicting non-essential genes is intrinsically more challenging than essential ones [4]. Our approach illustrates the potential of exploring new ways of combining traditional tools such as Flux Balance Analysis with modern data-driven approaches, and adds to the growing body of literature at the interface of genome-scale metabolic modeling with machine learning [40, 47].

## IV METHODS

### A Flux Balance Analysis

Flux Balance Analysis (FBA) is one a popular methods for the analysis of cellular metabolism. In steady state, a metabolic network can be described by

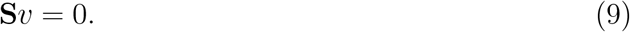

The aim of FBA is to obtain the solution vector *v*^∗^ that satisfies the above condition and at the same time solves the following optimization problem:

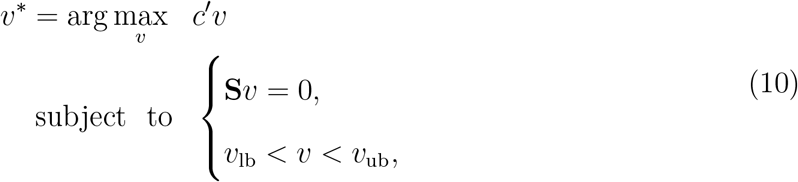

in which, *c* is a vector of flux weights, and (*v*_lb_, *v*_ub_) are lower and upper bounds on reaction fluxes, respectively.

### B Mass Flow Graphs

Originally introduced in [3], MFGs are designed to reflect the the directional flow of metabolites produced or consumed through enzymatic reactions. In these graphs, reactions are considered as vertices and two reactions are connected through a directed edge is they share a metabolite (either as reactants or products). The construction pipeline of these graphs can incorporate different experimental conditions through varying flux distributions. For instance, one key advantage of such pipeline is the automatic pruning of pool metabolites through mapping onto weak connections between graph nodes, and therefore reducing their impact on the overall network structure.

To construct an MFG from a metabolite network consisting of *m* reactions and *n* metabolites, first we obtain the solution vector *v*^∗^ from FBA. Then, we unfold the *v*^∗^ into two fold forward and reverse reaction fluxes through

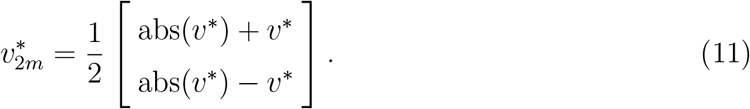

Next, the corresponding stoichiometric matrix of 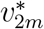 is defined as

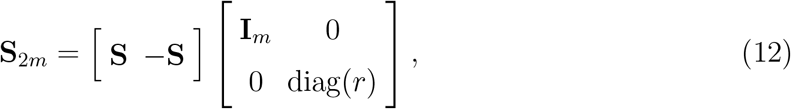

in which, S in the *n×m* stoichiometric matrix corresponds to *n* reactions and *m* metabolites of the original network, and *r* is an *m* dimensional Boolean vector indicating whether a reaction is reversible or not. Finally, the adjacency matrix of the MFG can be calculated as

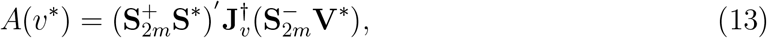

where † is the matrix pseudoinverse operator, and 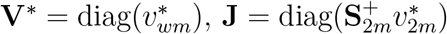 with

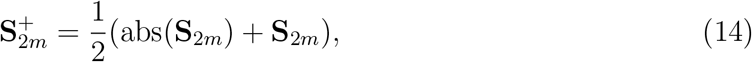

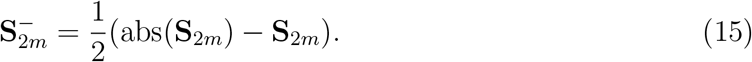

### C Node feature generation

Mass Flow Graphs are none-attributed, i.e., no specific node features are provided for reactions (nodes) in the graph. As a result, it is necessary to design a feature generation pipeline that considers the structure of the graph as well as the edge weights that appear in the adjacency of the graph. For this task, we propose a node encoding algorithm analogous to positional encoding of Transformer architectures.

Apart from the proposed FPE features in Eq. (6), a second approach to node encoding is to gather local neighbourhood structural statistics based on degree of each node and its neighboring nodes [7]. In this approach, the local degree profile of each node is defined as

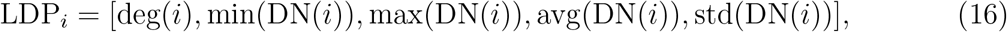

in which DN(*·*) is the operator for the set of neighboring nodes out-degree values. Additionally, a third encoding method relies on random walks from each node. In this method, a random walk encoding for each node *i* is calculated as:

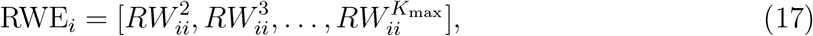

where *RW* = **AD**^−1^ is the random walk operator and only the random walks that end in node *i* are considered.

### D Message-passing neural networks (MPNN)

For representation learning of the graph features we employed GAT architecture [46] which is an instance of a MPNN scheme. In a typical graph representation learning task, the representation of each node is updated through a message passing scheme in which the information from neighboring nodes are gathered using message formula and aggregated with the features of the node itself. Thus, a message passing formula for each message from node *j* to node *i* can be written as

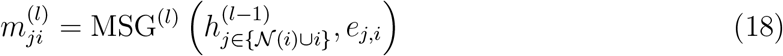

where 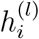 is the representation vector of node *i* in layer (*l*) of the MPNN, *e*_*j,i*_ are the features of the edge between node *i* and node *j*, and 𝒩 (*i*) is the set of neighbouring nodes to node *i*. Moreover, the messages for each node are aggregated to obtain the representation of node *i* in layer *l* using

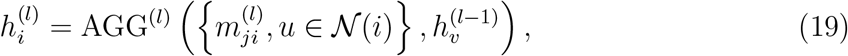

in which, AGG is a custom permutation invariant operator with regards to messages for each node.

GAT formulates the message equation in (18) as the multiplication of the attention as the learnable importance factor of each message by the representation of neighbours. Thus, the formula in (18) becomes:

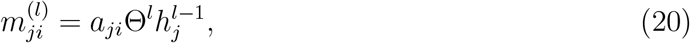

in which, *a*_*ji*_ is the attention coefficient and is usually calculated through feeding the features of the both neighbouring nodes *i* and *j* through a learnable function and calculating the importance through softmax function. Other popular examples of MPNN framework are Graph Convolution Network (GCN) [25] and GraphSAGE [20] which change the message function and use different aggregation functions. In GCN, the message function in (18) is calculated as:

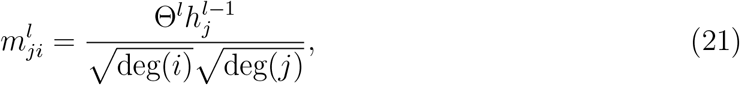

with the sum pooling operator as the aggregator function. In GraphSAGE, the message function MSG is the identity function and the aggregation function AGG is calculated as:

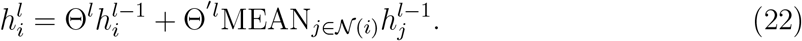

where MEAN is the mean pooling operator. The comparison between the performance of different MPNN schemes is presented in the Supplementary Figure S1.

### E Graph construction

To build the MFGs, we employed the iML1515 model of *E. coli* MG1655 introduced by Monk *et al* [31]. To label the reaction nodes in the graph, we employed the growth assay data from the same work on strain BW25113. Since BW25113 lacks several genes from MG1655, we produced FBA solutions by setting their reaction bounds to zero and assuming aerobic growth. The reaction bounds can be found in Supplementary Table S1. To simulate *E. coli* growth in a specific carbon sources, we set the corresponding exchange flux to a fixed value and deactivate all other carbon exchange fluxes. The list of all carbon sources and their corresponding exchange reactions can be found in Supplementary Table S2. All calculations were done with the COBRApy toolbox v0.26.3 using the glpk solver and the default objective function included in the iML1515 model.

### F Performance evaluation of binary classifiers

#### a. Training and evaluation in a single carbon source

We start our evaluations of FlowGAT from MFG resulting from glucose as the sole carbon source in Figure 2A. After mapping reactions to genes based on GPR rule set, growth rate values were converted to essentiality labels based on the threshold of 0.5 and were assigned to corresponding nodes in the MFG. As mentioned in Section II A, the labeled nodes in the MFG graph are imbalanced with a higher number of essentials compared to non-essential nodes. Therefore, for model training, we employed stratified sampling into 5 folds using built-in scikit-learn [5] functions with 1 fold for testing and 4 folds for training with the labeled nodes and 25% of the training set is set chosen as validation set (Figure 1D). For the initial tuning of hyperparameters, we employed grid search for each model and chose the best model settings based on the performance on the validation set; hyperparameters were kept constant for all other evaluations in the paper. All GNN based models were implemented and trained using the GraphGym in PyG package [14]; classic models (SVC, MLP, RF) were implemented using scikit-learn. The list of chosen hyperparameters for each model is available in Supplementary Table S3.

Due to the small number of available labeled data in our dataset (255 in case of glucose MFG), to compare the performance of different mmodels (Figure 2) we trained the GNN models on the training folds and evaluated the performance on the test fold 5 times, each time changing the train and test fold to ensure that the results are not caused by split bias. In the case of GNN based models, for each evaluation step, 25% of the training fold was considered as the early stopping set. We kept track of the best model on the early stopping set, in terms of the loss value after each training epoch, until the maximum number of epochs was reached. The weights of the best model at the end of training were then saved and employed to predict for the nodes of the test fold. Additionally, each training and evaluation step on a test fold was repeated 10 times with the model retrained with a different initial random seed to make sure the predictions were not a result of random seed selection for weight initialization. In total, 50 evaluation steps (5 folds and 10 times for each fold) were gathered for each model. For the classic models (SVC, RF, MLP), we employed the same process except for the use early stopping set. We followed the same procedure for model evaluation in other carbon sources (Figure 3B).

#### b. Training and evaluations across carbon sources

To produce the evaluations in Figure 3C, for each MFG the training and early stopping folds were chosen with a 4:1 ratio; in all cases we tested each model on all nodes of the other MFGs. Following the same scheme as in the previous section, the best performing model on the early stopping set was chosen for the evaluation of the test set; the training set was resampled 5 times and each model was retrained 10 times with different initial weights.

In Figures 3B–C, we quantified the performance improvement of FlowGAT over the no-skill classifier 100 *×* (PRAUC_FlowGAT_ − PRAUC_no-skill_) */*PRAUC_no-skill_.

## ACKNOWLEDGMENTS

TM was supported by the Research Council of Norway (grant number 312045). DAO was supported by the United Kingdom Research and Innovation (grant EP/S02431X/1, UKRI Centre for Doctoral Training in Biomedical AI. Computations were performed on resources provided by Sigma2 - the National Infrastructure for High Performance Computing and Data Storage in Norway (project NS9715).

## Supplementary Figures and Tables

**Supplementary Figure S1.**
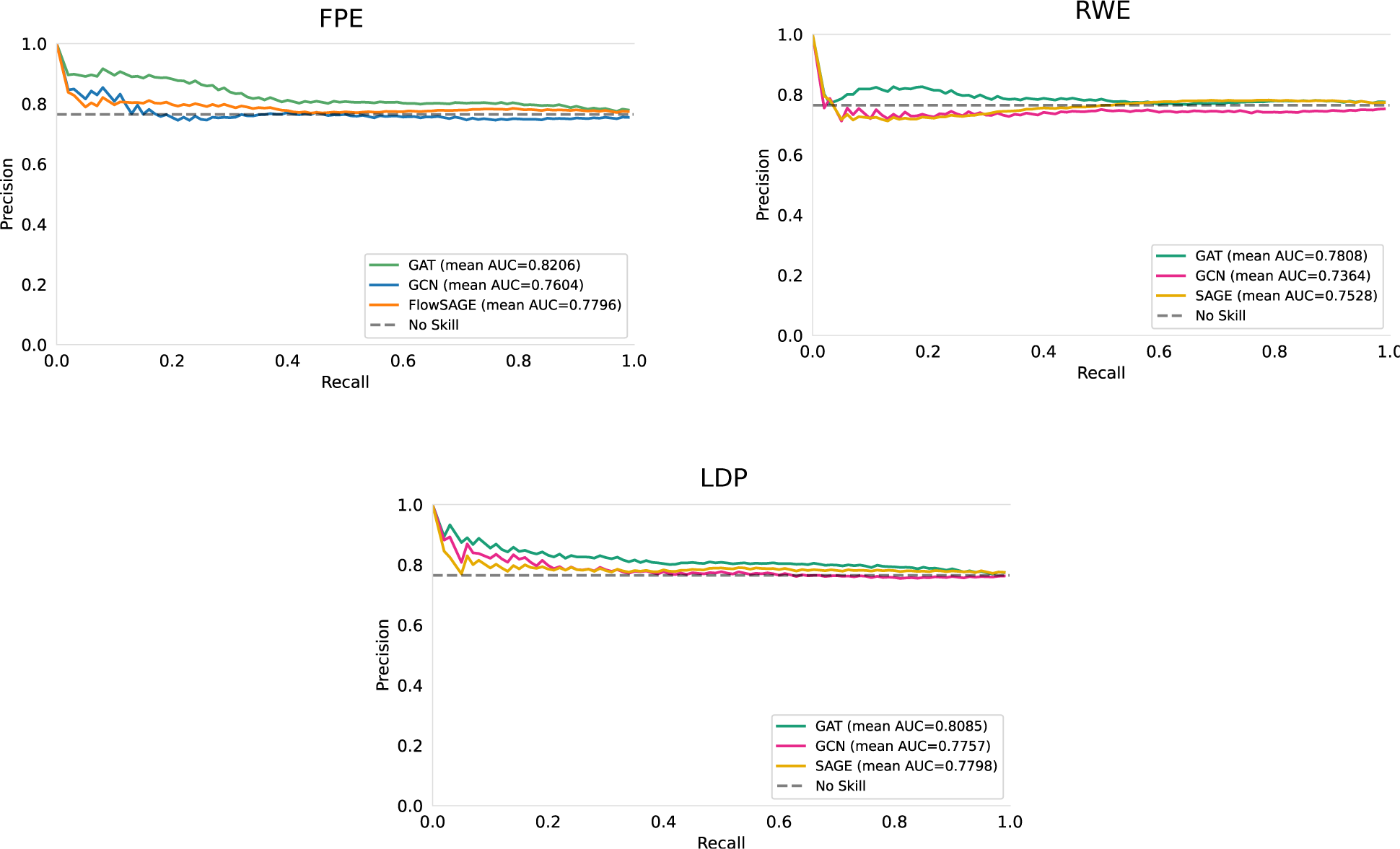
Performance comparison between different message passing schemes and node embeddings. Model training and evaluation followed the same procedure as Figure 2A in the main text. Results show three node embeddings (Flow Profile Embeddings defined in Eq. (6), Local Degree Profile, and Random Walk Embeddings as explained in the Methods) and three popular message passing schemes (Graph Attention as in Figure 2A, Graph Convolution Networks, and Graph SAGE).

**Supplementary Figure S2.**
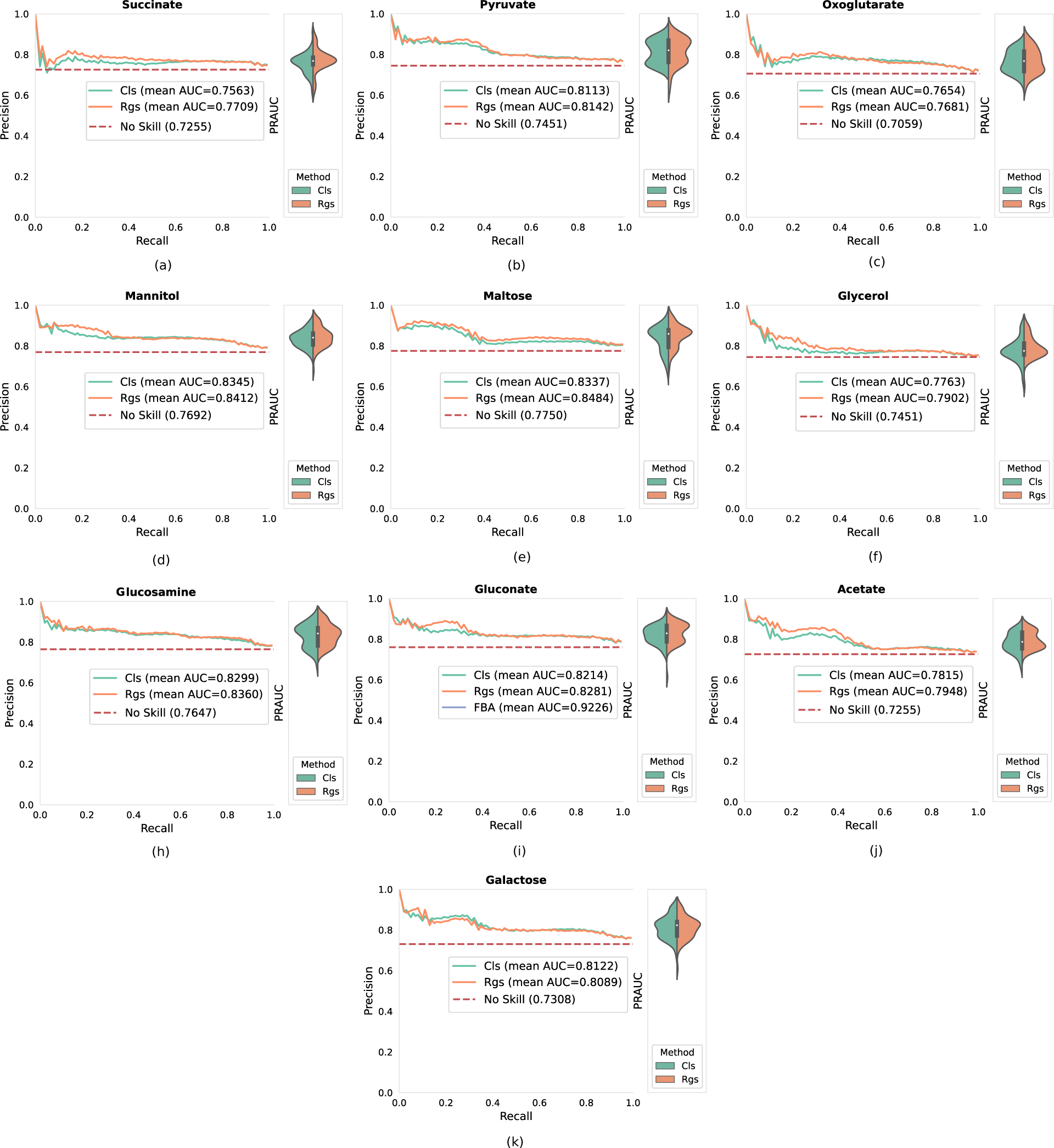
Performance comparison of classification and regression training for FlowGAT trained on labels and mass flow graphs for various carbon sources. For each of the ten graphs, FlowGAT was trained on the nodes of the graph using both classification and regression training scheme, as in Figure 2C in the main text. Precision-recall curves were averaged over 5 cross-validation folds and 10 rounds of re-training for different random seeds.

**Supplementary Table S1.** Bounds for exchange reactions for the *E. coli* iML1515 model growing in aerobic conditions with glucose as the only carbon source.

| Reaction | Bounds |
| --- | --- |
| L-arabinose isomerase (ARAI) | $v_{lb} = v_{ub} = 0$ |
| L-ribulokinase (RBK_L1) |  |
| Rhamnulose-1-phosphate aldolase (RMPA) |  |
| Lyxose isomerase (LYXI) |  |
| L-rhamnose isomerase (RMI), |  |
| Rhamnulokinase (RMK) |  |
| B-galactosidase (LACZ) |  |
| Oxygen exchange (EX_o2_e) | $v_{lb} = -20$ |

**Supplementary Table S2.**
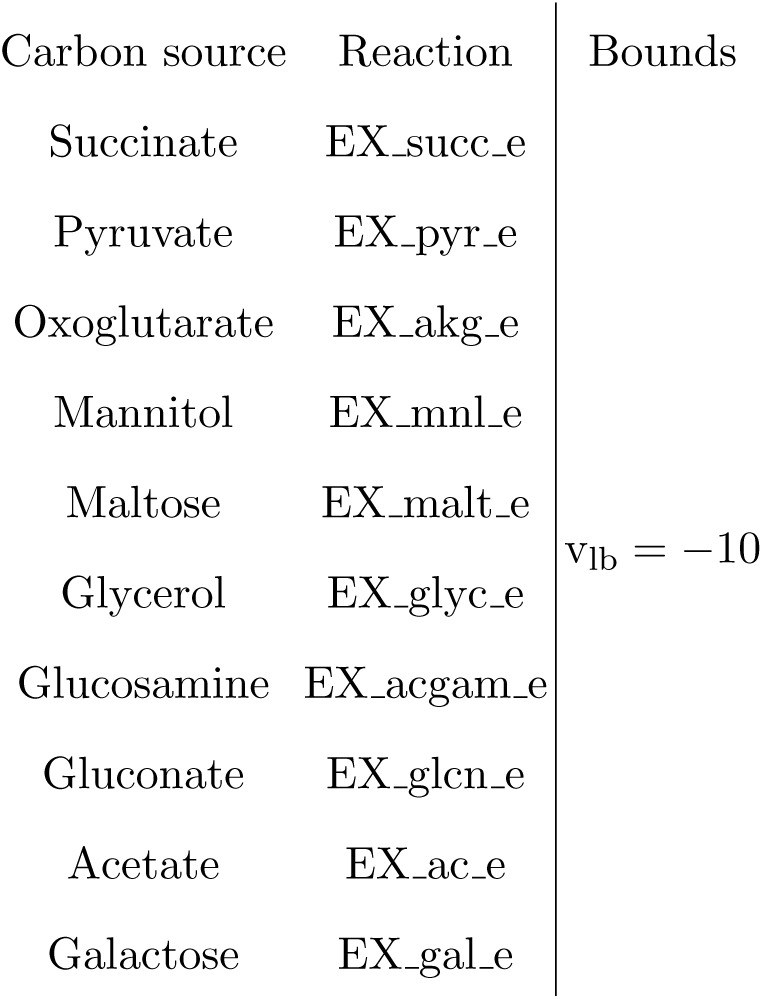
Exchange reactions for different carbon sources in the *E. coli* iML1515 genome-scale metabolic model. For each carbon source, the corresponding reaction bounds were adjusted and all other reaction bounds for other sources were set to 0.

**Supplementary Table S3.**
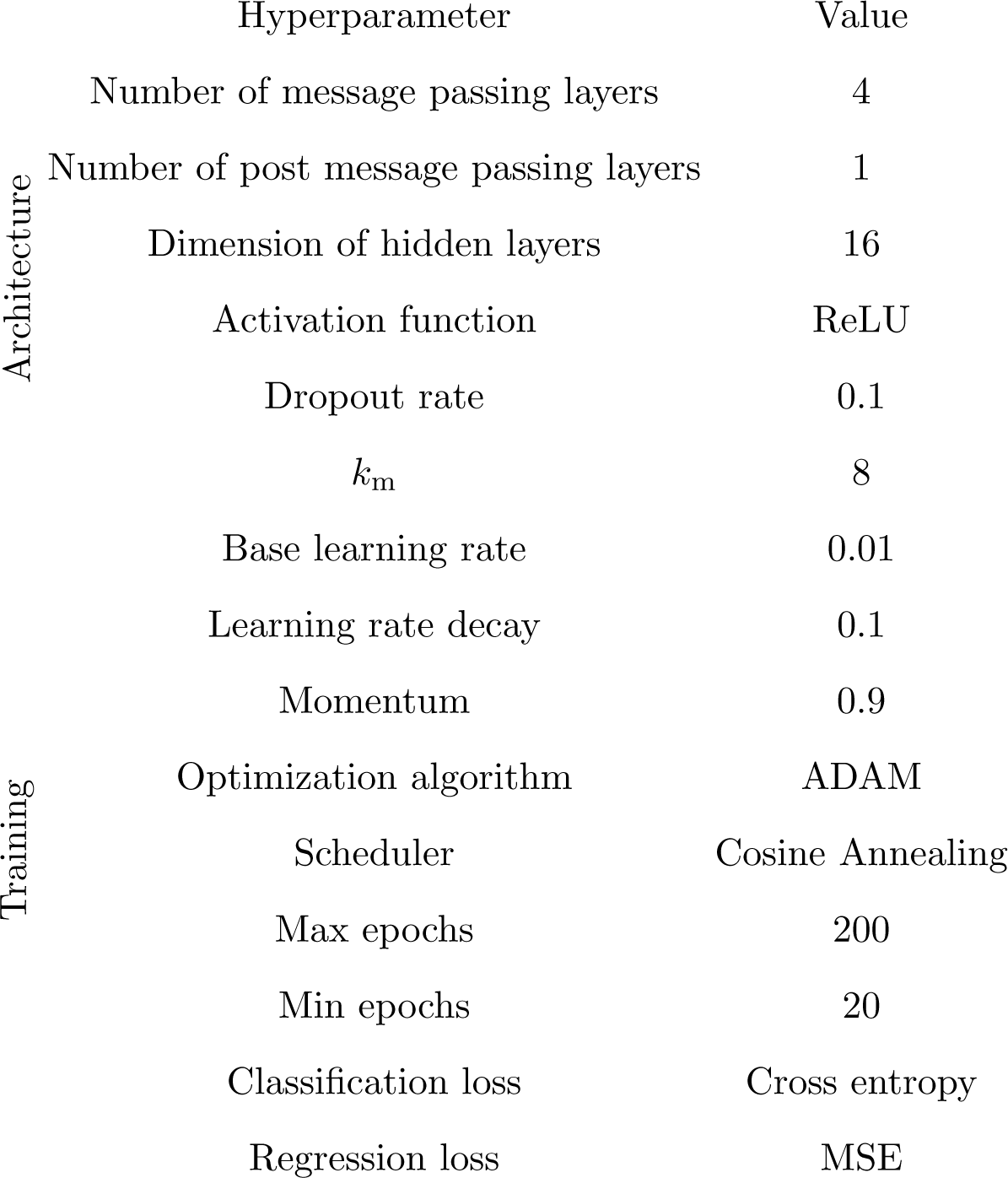
Hyperparameters list for GNN based models. For each GNN based model the list of above hyperparameters was determined after initial tuning on a chosen validation set through grid search. Afterward, the above set was used for all evaluations reported in the paper. For a fair comparisons amongst different GNNs, the architecture and training hyperparameters are kept the same and the message passing formula is the only change in the architecture (e.g., GAT, GCN, SAGE)

